# Periosteum-derived Osteocrin regulates bone growth through both endochondral ossification and intramembranous ossification

**DOI:** 10.1101/697094

**Authors:** Haruko Watanabe-Takano, Hiroki Ochi, Ayano Chiba, Ayaka Matsuo, Yugo Kanai, Shigetomo Fukuhara, Keisuke Sako, Takahiro Miyazaki, Shingo Sato, Naoto Minamino, Shu Takeda, Akihiro Yasoda, Naoki Mochizuki

## Abstract

During development of long bones, two mechanistically distinct processes contribute to long- and short-axis growth. Endochondral ossification in the growth plate leads to the long-axis growth, while intramembranous ossification including apposition in the periosteum regulates the short axis growth. Here, we show that periosteal osteoblast-derived secretory peptide, Osteocrin (OSTN), promotes both types of long bone growth through potentiation of signaling by C-type natriuretic peptide (CNP), because OSTN inhibits the clearance of CNP by binding to natriuretic peptide receptor 3 (NPR3). The mice lacking OSTN showed less bone mass in trabecular and cortical regions than the control mice, suggesting the dual functions of OSTN in long bone growth. We found that OSTN regulated trabecular bone formation by inducing proliferation and maturation of chondrocytes possibly through enhancing CNP-dependent signaling. Besides the contribution of OSTN to long axis growth, we demonstrated that OSTN together with CNP induced osteoblast differentiation of periosteum-derived multipotent progenitor cells expressing NPR3. These data suggest that OSTN induces long bone growth through endochondral ossification and osteoblast specification of multipotent progenitor cells in the periosteum.

## INTRODUCTION

The long bones develop through two types of ossifications: endochondral ossification and intramembranous ossification, which regulate growth in length and width of long bone, respectively. Endochondral ossification is the process by which chondrocytes of the cartilage template are replaced by bone (trabecular bone), while intramembranous ossification is that by which osteoblasts directly produce mineralized bone in the inner layer of periosteum (cambium layer) (Kozhemyakina et al., 2015; Long and Ornitz, 2013; Mackie et al., 2011; Rauch, 2005).

During endochondral ossification, chondrocytes are organized into the growth plate located between the epiphysis and metaphysis and undergo a sequential maturation process to become hypertrophic chondrocytes that are ultimately replaced by bone. Several secretory factors such as parathyroid hormone-related protein (PTHrP), Wnt ligands, Sonic hedgehog (SHH), Indian hedgehog (IHH), and fibroblast growth factor (FGF) mediate endochondral ossification (Kozhemyakina et al., 2015; Long and Ornitz, 2013; Mackie et al., 2011). In addition, C-type natriuretic peptide (CNP), a member of natriuretic factors (NPs), regulates endochondral ossification by promoting chondrocyte proliferation and maturation. CNP stimulates its receptor guanylate cyclase (GC)-B to produce cyclic GMP (cGMP), leading to activation of cGMP-dependent protein kinases (cGKs) (Potter et al., 2006). Previous studies have shown that the mice lacking CNP, GC-B, or cGKII exhibited short stature, while overexpression of CNP in mice resulted in skeletal overgrowth (Chusho et al., 2001; Kawasaki et al., 2008; Nakao et al., 2015; Tsuji and Kunieda, 2005; Yasoda et al., 2004). These findings suggest an essential role for a CNP-GC-B-cGKII signaling pathway in endochondral ossification. In addition, the mice deficient for natriuretic peptide receptor 3 (NPR3), a clearance receptor for NPs, showed bone elongation, suggesting that proper formation of long bone requires tight regulation of CNP signaling by NPR3-mediated clearance of CNP (Jaubert et al., 1999; Matsukawa et al., 1999).

During intramembranous ossification, skeletal stem cells in the periosteum contribute to appositional growth by becoming bone-forming osteoblasts to increase bone thickness and to heal fracture (Debnath et al., 2018; Duchamp de Lageneste et al., 2018). Clinical observations using healthy human subjects reveal spatiotemporal variability of the rate of appositional growth and gender difference in bone growth (Baxter-Jones et al., 2011; Gabel et al., 2015). Moreover, it has been reported that differentiation and proliferation of periosteal osteoblast progenitors are regulated by estrogen and parathyroid hormone (PTH), suggesting the regulation of intramembranous ossification by systemic factors (Ogita et al., 2008). In addition, expression analysis of periosteum-derived cells reveals that intramembranous ossification in periosteum is regulated by local factors such as Wnt, hedgehog (Hh), and chemokines (Kim et al., 2007; Stich et al., 2008; Wang et al., 2010). Thus, differentiation of skeletal stem cells in the periosteum into osteoblasts might be regulated by both systemic and local cues. However, little is known about the mechanism by which those stem cells differentiate into osteoblasts to facilitate appositional growth.

*Osteocrin* (OSTN), also known as *Musclin,* encodes a secretory protein that is expressed in bone and skeletal muscles (Nishizawa et al., 2004; Thomas et al., 2003). OSTN contains two regions with high homology to NPs. Although OSTN does not bind to GC-A and GC-B due to the lack of conserved cysteine ring (Moffatt et al., 2007), it specifically binds to NPR3 with similar affinity to NPs (Kita et al., 2009). Accordingly, OSTN competes with NPs for binding to NPR3, thereby increases NPs availability by protecting them from NPR3-depenedent degradation (Kanai et al., 2017; Kita et al., 2009; Miyazaki et al., 2018; Moffatt et al., 2007; Subbotina et al., 2015). Previously, we and others have reported that overexpression of OSTN in transgenic mice induces an increase in bone length by enhancing CNP signal-mediated proliferation and maturation of chondrocytes (Kanai et al., 2017; Moffatt et al., 2007). In addition, our analysis of OSTN mutant zebrafish revealed that OSTN is also involved in bone and cartilage development in zebrafish (Chiba et al., 2017). However, the role of endogenous OSTN in mammalian bone development and the mechanism by which OSTN regulates bone growth are still obscure.

In this study, we analyzed OSTN-deficient mice and found that endogenous OSTN regulates endochondral ossification during long bone formation by inducing proliferation and maturation of chondrocytes possibly through enhancing CNP signaling. In addition, we showed that OSTN promotes cortical bone formation by facilitating CNP signal-mediated differentiation of osteoblasts and unveiled the role of OSTN and CNP in intramembranous ossification. We, therefore, propose that OSTN potentiates long bone growth through promoting both endochondral ossification and intramembranous ossification by regulating CNP signaling.

## RESULTS

### OSTN is expressed in periosteal osteoblasts during bone development

To examine the distribution of OSTN, we generated knock-in mice in which a LacZ reporter gene and neomycin cassette are inserted upstream of exon 3 of *Ostn* using the targeting vector obtained from Knockout Mouse Project (KOMP) repository (Fig. S1A). We used heterozygous mice (*Ostn*^+/LacZ^) to detect LacZ activity as an indicator of *Ostn* promoter activity. X-gal staining in whole cleared body revealed that *Ostn* expression in some bones started at embryonic 15.5 (E15.5). The expression became more appreciable at the age of 30 days (P30) (Fig. 1A). *Ostn* expression was evident in long bones (zygomatic bones, mandible, tibias, metacarpal bones, phalanges, caudal vertebrae, ulnae, radii, and ribs) as well as flat bones (nasal bones, frontal bones, parietal bones, occipital bones, and scapulas), but faint in vertebrae, femora, and humeri. At the age of 3 months, *Ostn* expression seemed to be entirely downregulated even in the bones in which *Ostn* expression was apparent at age 30 days.

**Fig. 1.**
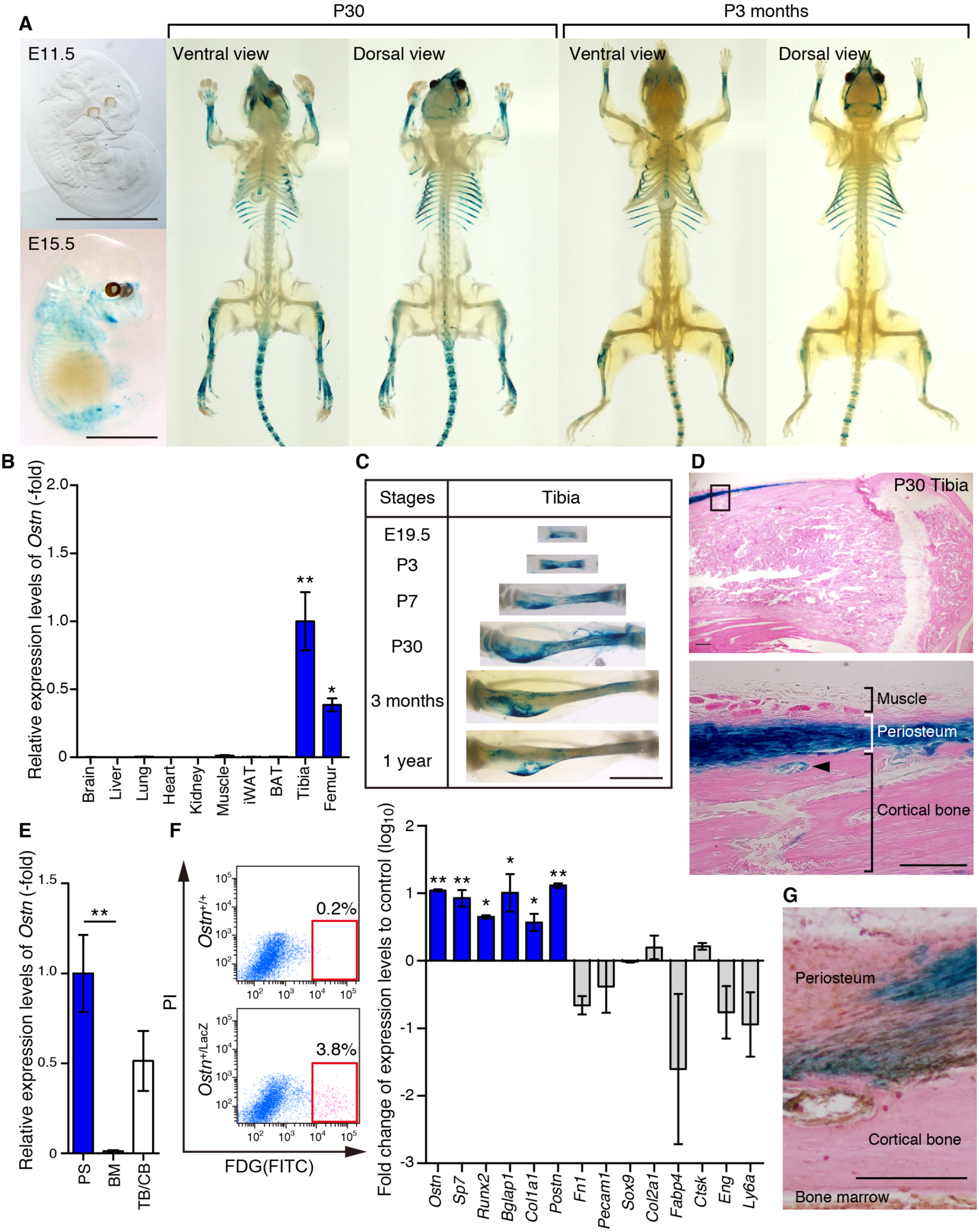
OSTN is expressed in periosteal osteoblasts during bone development. (A) LacZ staining of cleared whole body of *Ostn*^+/LacZ^ mice at the stage indicated at the top. E, embryonic day; P, postnatal. (B) *Ostn* mRNA expression in various tissues of 8-week-old ICR male mice. Gene expression value of each sample was normalized by each *Gapdh* mRNA value. The relative expression to the value of tibia is indicated. (n=4) (C) LacZ staining of cleared tibia of *Ostn*^+/LacZ^ mouse at the stage indicated by the left. (D) Eosin and LacZ staining of tibia of the *Ostn*^+/LacZ^ mice at the age of P30. Note that OSTN is expressed in the periosteum but not in growth plate and endosteum. Arrow head indicates basic multicellular unit in compact bone. (E) Relative expression levels of *Ostn* in fractionated bone; periosteum (PS), bone marrow (BM), and trabecular/cortical bone (TB/CB). (n=4) (F) Representative plot images of flow cytometry (left). x and y axis represent fluorescein di-D-galactopyranoside (FDG)-derived FITC and Propidium iodide (PI), respectively. Note the presence of LacZ-positive (red box) population in *Ostn*^+/LacZ^ mouse, but not in *Ostn*^+/+^ mouse. Marker analysis of LacZ-expressing cells by qPCR (right). Values represent log10 fold-change of the expression levels in LacZ-positive cells relative to that in LacZ-negative cells. (n=3) (G) Eosin staining with LacZ staining (blue) and immunostaining using Osterix antibody (brown) of the tibia of the *Ostn*^+/LacZ^ mice at the age of P30. Scale bars: A,C, 0.5 cm; D,G, 200 μm. Data are mean ± s.e.m. ** *P* < 0.01, * *P*<0.05; by One-way ANOVA followed by Turkey-Kramer test in B, E and by Student’s t test in F.

To confirm whether LacZ-positive staining reflects the endogenous *Ostn* expression, we conducted qPCR analyses using the tissues isolated from 8-week-old mice. Consistent with the results of the LacZ staining, *Ostn* expression in tibia and femur was significantly higher than that in other tissues (Fig. 1B, Fig. S1B). X-gal staining images of tibia at the various developmental stages demonstrated that *Ostn* expression increased until the age of 30 days and decreased by aging, suggesting its biological significance in bone morphogenesis (Fig. 1C). To examine where *Ostn*-expressing cells reside in tibia, we histologically examined the expression in X-gal-stained sections (Fig. 1D). It is of note that the expression of *Ostn* was high in cells of periosteum and detectable in basic multicellular unit in compact bone (arrowhead). On the other hand, any staining of *Ostn* expression was not detected in growth plate and endosteum.

To further validate whether LacZ staining reflects the endogenous *Ostn* expression, we analyzed *Ostn* expression using the fractionated RNA samples of bones by qPCR. Successful fractionation was verified by the expression of specific marker genes (Fig. S1C). The expression levels of *Ostn* in periosteum (PS) and trabecular/cortical bone (TB/CB) were higher compared to that in bone marrow (BM) fraction (Fig. 1E). To characterize the *Ostn*-expressing cells, we isolated the cells exhibiting high LacZ activity by using fluorescence activated cell sorting (FACS) and analyzed the expression of marker genes specific for individual cell types by qPCR (Fig. 1F). We found that *Ostn*-expressing cells belong to certain population of periosteum-derived cells and that specific markers for osteoblasts were highly expressed in *Ostn*-expressing cells. In contrast, the expression of markers for other cell types, including fibroblasts, endothelial cells, chondrocytes, adipocytes, osteoclasts, and mesenchymal cells, were low or comparable in LacZ-positive cells compared with that in LacZ-negative cells. Furthermore, immunostaining with the antibody for Osterix, a specific marker for osteoblast, revealed that the LacZ-positive cells partially overlapped with Osterix-positive cells (Fig. 1G). These results suggest that not all but certain osteoblasts in the periosteum express OSTN.

### OSTN-deficient mice exhibit mild growth retardation and skeletal abnormality

To elucidate physiological roles of OSTN, we generated homozygous mutant mice (*Ostn* ^LacZ/LacZ^ mice) harboring LacZ and neomycin inserts in exon 3 of *Ostn* gene locus as shown in Fig. S1A. qPCR analysis revealed significant reduction of *Ostn* transcripts in *Ostn* ^LacZ/LacZ^ mice as compared with that in wild-type mice (*Ostn*^+/+^ mice) (Fig. 2A). Thus, we used these mice as *Ostn*-deficient mice*. Ostn*^LacZ/LacZ^ mice were fertile and had normal gross appearance except for mild growth retardation (Fig. 2B). The body weight and body length of *Ostn*^LacZ/LacZ^ mice at the age of postnatal 8 weeks were decreased as compared to the controls (Fig. 2C,D). On the other hand, the ratios of tissue weight to body weight of liver, heart, kidney, and lung were comparable to that in *Ostn*^+/+^ mice (Fig. 2E). Whole-body imaging of skeletal systems scanned by computed tomography (CT) revealed slight shortening of some bones in 8-week-old *Ostn*^LacZ/LacZ^ mice (Fig. 2F). The length of femora, tibiae, and caudal vertebrae except for carvarie of *Ostn*^LacZ/LacZ^ mice at the age of postnatal 8 weeks was significantly shorter than that of *Ostn*^+/+^ mice (Fig. 2G). These results suggest that OSTN is required for normal skeletal development of long bones.

**Fig. 2.**
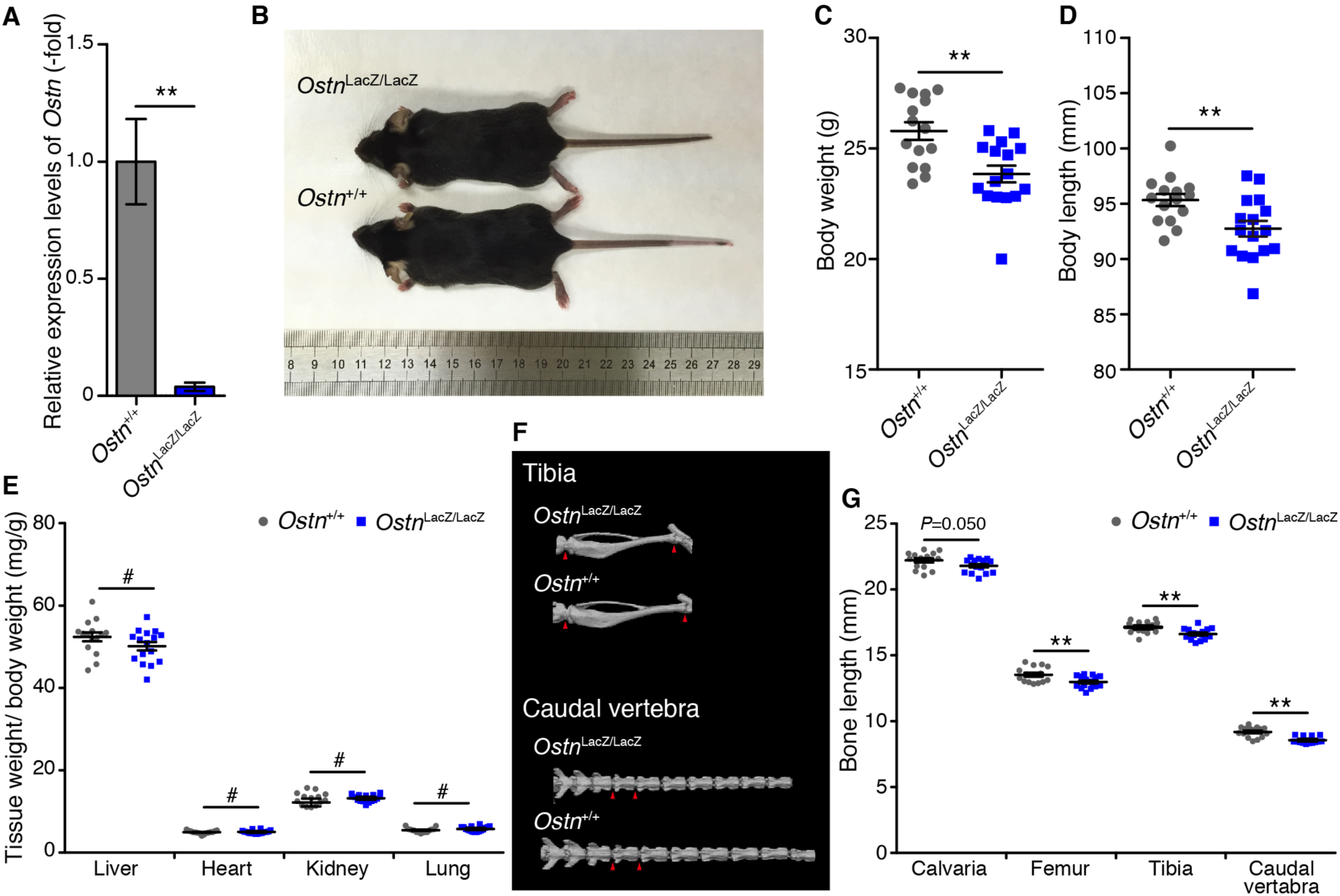
OSTN-deficient mice exhibit mild growth retardation and skeletal abnormality. (A) qPCR analysis of *Ostn* expression in the hind limbs of *Ostn*^LacZ/LacZ^ and *Ostn*^+/+^. Relative expression to the value obtained from *Ostn*^+/+^ is indicated similarly to Fig. 1B. (n=5) (B) Gross body appearance of male 8-week-old *Ostn*^LacZ/LacZ^ and *Ostn*^+/+^. Note that mild growth retardation of *Ostn*^LacZ/LacZ^ mouse. Body weight (C) and body length (D) of 8-week-old *Ostn*^LacZ/LacZ^ mice (n=16) and *Ostn*^+/+^ mice. (n=15) (E) Comparison of normalized tissue weights by body weights between of *Ostn*^LacZ/LacZ^ and *Ostn*^+/+^ mice. (n=15-16) (F) Computed tomography (CT) images of tibiae (upper) and caudal vertebrae (lower) in *Ostn*^LacZ/LacZ^ and *Ostn*^+/+^. Red arrow heads indicate ends of each bone. (G) Comparison of bone (as indicted at the bottom) length between *Ostn*^LacZ/LacZ^ and *Ostn*^+/+^ mice. Note the shortened bone length of *Ostn*^LacZ/LacZ^. Data are mean ± s.e.m. (n=15-16) ** *P* < 0.01, * *P* <0.05, # *P* ≥0.05 by Student’s t test.

### Impairment of growth plate growth and trabecular bone in OSTN-deficient mice

It has been known that a CNP-GC-B-cGKII axis plays a pivotal role in morphogenesis of growth plate, which extends long bones (Chusho et al., 2001; Kawasaki et al., 2008; Nakao et al., 2015; Tsuji and Kunieda, 2005; Yasoda et al., 2004). Consistently, NPR3-deficient mice exhibited skeletal overgrowth with elongated growth plate (Matsukawa et al., 1999). We have shown that transgenic mice over-expressing OSTN show skeletal overgrowth by inhibiting the NPR3-dependent degradative pathway of NPs (Kanai et al., 2017). Therefore, we assumed that OSTN has a strong relevance to endochondral ossification. To elucidate whether endogenous OSTN is required for morphogenesis of growth plate, we first histologically analyzed the growth plates of *Ostn*^LacZ/LacZ^ mice at postnatal 3 and 8 weeks. As shown in Fig. 3A, *Ostn*^LacZ/LacZ^ mice had thinner growth plates than the control mice. Quantitative analyses indicated a significant reduction of growth plate thickness of *Ostn*^LacZ/LacZ^ mice as compared with *Ostn*^+/+^ mice at the ages of both 3 and 8 weeks (Fig. 3B).

**Fig. 3.**
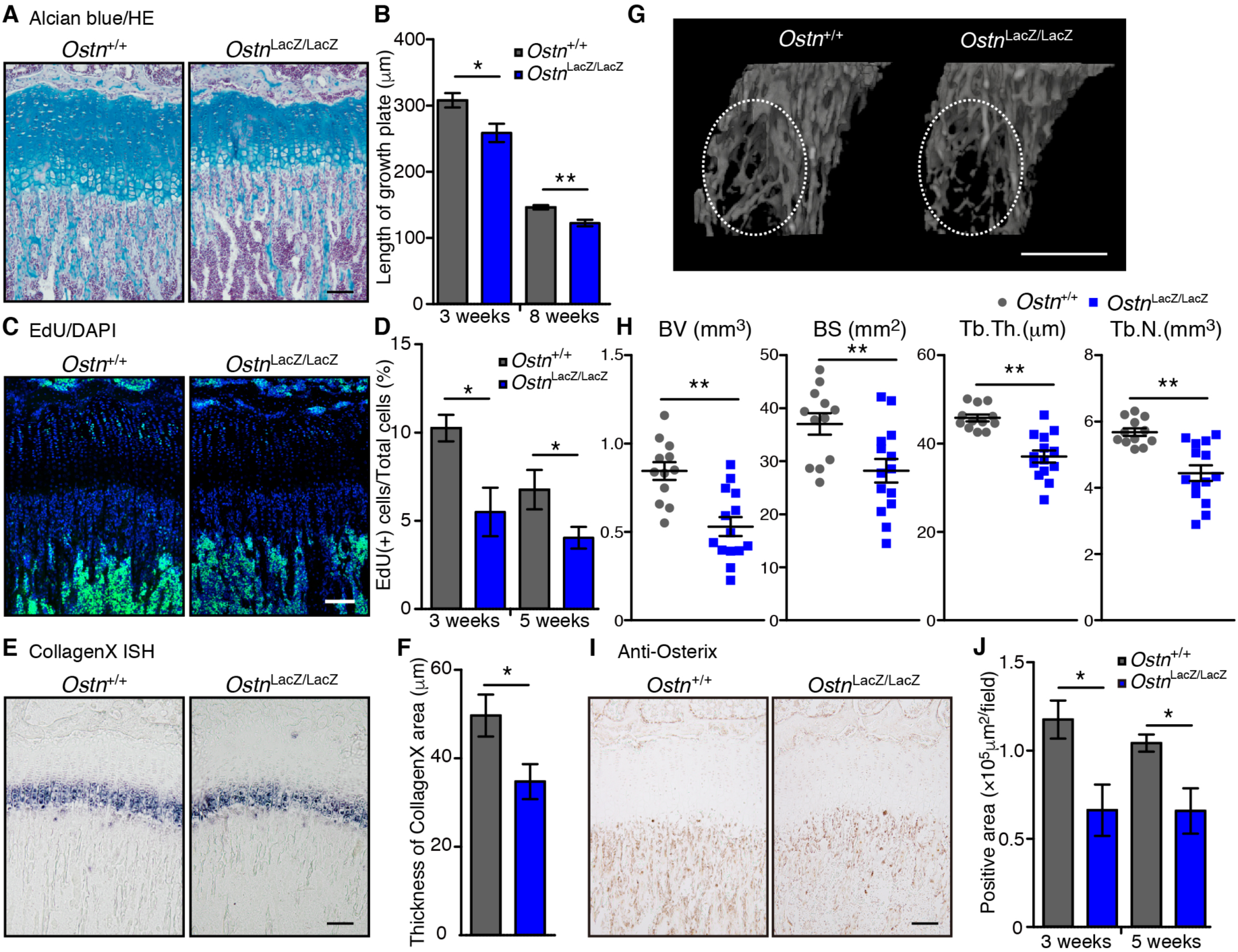
Impairment of growth plate growth and trabecular bone in OSTN-deficient mice. (A) Representative images of Alcian Blue and hematoxylin eosin staining of growth plates of the 3-week-old *Ostn*^LacZ/LacZ^ and *Ostn* ^+/+^ mice. (B) Thickness of growth plates of the 3- and 8-week-old *Ostn*^LacZ/LacZ^ and *Ostn* ^+/+^ mice. (n=6-9) (C) Representative images of staining of growth plates of 3-week-old *Ostn*^LacZ/LacZ^ and *Ostn* ^+/+^ mice. EdU-positive cells (green) and DAPI-positive nuclei of all cells (blue). (D) Percentages of the number of EdU-positive cells in that of total cells in growth plates of 3- and 5-week-old *Ostn*^LacZ/LacZ^ and *Ostn* ^+/+^ mice. (n=4-6) (E) Images of in situ hybridization for *Col10a1* of the growth plate of 5-week-old *Ostn*^LacZ/LacZ^ and *Ostn* ^+/+^ mice. (F) Quantitative analyses of the thickness of *Col10a1*-positive area in the sections used in (E). Note decreased maturation of chondrocytes in the *Ostn*^LacZ/LacZ^. (n=5) (G) Representative 3D μCT images of trabecular bone at proximal-metaphyseal side of tibia in the 8-week-old *Ostn*^LacZ/LacZ^ and *Ostn* ^+/+^ mice. (H) Measurement of trabecular bone parameters: bone volume (BV), bone surface (BS), trabecular thickness (Tb.Th.), and trabecular number (Tb.N.) assessed by μCT. (n=12-14) (I) Representative images of immunostaining of growth plates of 5-week-old *Ostn*^LacZ/LacZ^ and *Ostn* ^+/+^ mice using anti-Osterix antibody, a marker of pre-osteoblasts. (J) The area of Osterix-positive cells in the growth plate of 3- and 5-week-old *Ostn*^LacZ/LacZ^ and *Ostn* ^+/+^ mice. (n=5) Data are mean ± s.e.m. \**P*<0.05, \*\**P*<0.01, by Student’s *t*-test. Scale bars: A,C,E,100 μm; G,1000 μm; I,50 μm.

Because an activation of CNP-GC-B-cGKII signaling pathway results in proliferation and maturation of chondrocytes in growth plates (Miyazawa et al., 2002; Nakao et al., 2015), we next examined the proliferation of chondrocytes in vivo by incorporation of EdU in vivo (Fig. 3C). Notably, the number of EdU-positive cells of *Ostn*^LacZ/LacZ^ mice was lower than that of the *Ostn*^+/+^ mice. We also noticed that the number of EdU-positive cells of both mice decreased gradually along with the progression of skeletal development (Fig. 3D). Furthermore, to evaluate chondrocytes maturation in the growth plate, Collagen, type X, alpha1 (*Col10a1*) expression was assessed by in situ hybridization (ISH) (Fig. 3E). We measured thicknesses of the area where *Col10a1*-expressing cells were detected and found a significant reduction of the thickness in *Ostn*^LacZ/LacZ^ mice as compared with *Ostn*^+/+^ mice at the age of 5 weeks (Fig. 3F). These results suggest that endogenous OSTN is required for chondrocyte proliferation and maturation in the growth plate, thereby contributing to skeletal growth.

Chondrocyte proliferation and maturation in the growth plate ultimately lead to endochondral ossification. To examine the consequence of impaired chondrogenesis of *Ostn*^LacZ/LacZ^ mice in the following endochondral ossification, we quantified bone microstructure of tibia using μCT. We observed a marked deficiency of microarchitecture of trabecular bones at proximal metaphysis of tibia in *Ostn*^LacZ/LacZ^ mice as compared with the controls at the age of 8 weeks (Fig. 3G,H, Fig. S2A). Although total bone volume (TV) of *Ostn*^LacZ/LacZ^ was comparable to that of *Ostn*^+/+^ mice, bone volume (BV), surface (BS), trabecular bone thickness (Tb.Th.), number (Tb.N.), and space (Tb.Sp.) were significantly decreased in *Ostn* ^LacZ/LacZ^ mice, indicating the failure of endochondral ossification in *Ostn* ^LacZ/LacZ^ mice. We further performed bone morphometric analyses and found reduced new bone formation in *Ostn* ^LacZ/LacZ^ mice as compared to *Ostn*^+/+^ mice, implying the impairment of osteoblast development in *Ostn*^LacZ/LacZ^ mice (Fig. S2B). To confirm this impairment in *Ostn*^LacZ/LacZ^ mice, we immunohistochemically analyzed Osterix expression (Fig. 3I). The staining of Osterix was significantly attenuated beneath the growth plate in *Ostn*^LacZ/LacZ^ mice (Fig. 3J). These results suggest that OSTN is required for endochondral ossification of tibia.

### OSTN-deficient mice show reduced cortical bone mass

To examine the effect of OSTN on short axis growth of bones, we imaged cross sectional area of tibia. Notably, growth of cortical bones in *Ostn*^LacZ/LacZ^ mice was reduced (Fig. 4A). μCT analyses of tibia revealed the thickness (Ct.Th.), area (Ct.Ar.), and other parameters (Tt.Ar. and Ct.Ar/Tt.Ar.) of cortical bones were attenuated in *Ostn*^LacZ/LacZ^ mice as compared with the controls (Fig. 4B), suggesting an unidentified OSTN-dependent mechanism that regulates cortical bone formation. There was a significant reduction in external line length (ELL), but not in internal line length (ILL) of *Ostn*^LacZ/LacZ^ mice, indicating the functional deterioration of osteoblast-dependent cortical bone formation in the outside of tibiae (Fig. 4C). To confirm the abnormality of osteoblast-dependent bone formation, we calculated the percentage of Osterix (+) cells isolated from the periosteum/perichondrium at the age of 11 days by using FACS (Fig. 4D,E). The percentages of Osterix (+) cells were significantly decreased in *Ostn*^LacZ/LacZ^ mice, suggesting the defective differentiation of osteoblasts from osteoblast progenitor cells in the periosteum in *Ostn*^LacZ/LacZ^ mice. These results at least suggest that OSTN is required for osteoblast-dependent cortical bone formation.

**Fig. 4.**
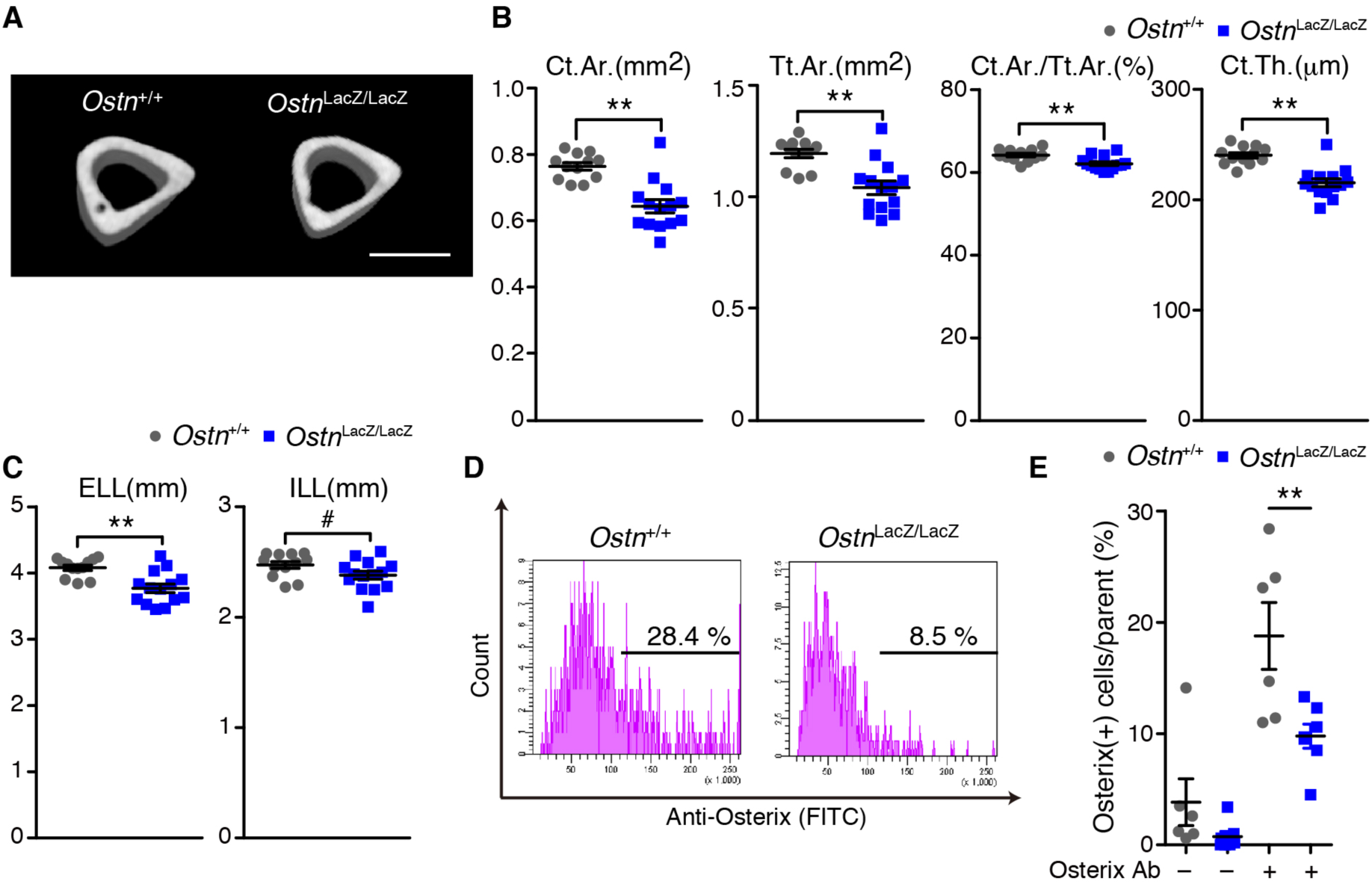
OSTN-deficient mice show reduced cortical bone mass. (A) Representative 3D μCT images of cortical bone at mid-diaphyseal side of tibia in 8-week-old *Ostn*^LacZ/LacZ^ and *Ostn* ^+/+^ mice. Scale bar; 1000 μm. (B,C) Measurement of cortical bone parameters: Cortical Area (Ct.Ar.), Total Area (Tt.Ar.), Cortical thickness (Ct.Th.), External line length (ELL), and Internal line length (ILL) (n=12-14). (D,E) Representative histograms of FACS analysis (D) and percentages of the number of Osterix-positive cells per parent population (E) using perichondrium of 11 days-old *Ostn*^LacZ/LacZ^ and *Ostn* ^+/+^ mice. Numbers in histograms indicate percentages of Osterix-positive cells per parent population. (n=6-7) Data are mean ± s.e.m. \**P*<0.05, \*\**P*<0.01, #*P*>0.05, by Student’s *t*-test.

### CNP promotes osteogenic differentiation of periosteum-derived cells

The effect of a CNP-GC-B axis on osteogenic differentiation from osteoblast progenitor cell is still obscure. Thus, we next wondered if CNP signaling pathway might regulate osteoblast-dependent cortical bone formation as well as endochondral ossification. We examined how CNP-dependent signaling contributes to osteogenic differentiation using periosteum-derived cells (PDCs) that contain specific skeletal stem cells for cortical bones (van Gastel et al., 2012).

First, we analyzed the expression of CNP (*Nppc*) and its receptors during osteogenic differentiation of PDCs. As shown in Fig. 5A, the expression of Osterix (*Sp7*) and Collagen, type I, alpha1 (*Col1a1*) of PDCs was gradually up-regulated in a time-dependent manner when cultured in the osteogenic differentiation medium (DM). Whereas the expression of *Npr3* was constant throughout the same period even though under differentiation condition, that of *Nppc* and GC-B *(Npr2*) was increased, suggesting a positive endogenous regulation of the CNP-GC-B axis for osteogenic differentiation. To investigate the potential of CNP signaling pathway in the osteogenic differentiation of PDCs, we examined the effect of exogenous CNP (CNP-22) on osteoblast differentiation. CNP-22 augmented alkaline phosphatase (ALP) activity and Ca^2+^ deposition stained with alizarin red S (ARS) in a dose-dependent manner under the differentiation conditions (Fig. 5B-D). The expression of *Sp7*, *Col1a1*, but not *Runx2* was augmented by CNP (Fig. 5E). In clear contrast, neither adipogenic differentiation nor chondrogenic differentiation from the PDCs examined by the oil red staining and micro mass culture followed by the staining with Alcian Blue, respectively was not augmented by CNP (Fig. 5F-I). Accordingly, these results suggest that CNP potentiates the specification and differentiation of PDCs into osteogenic lineage.

**Fig. 5.**
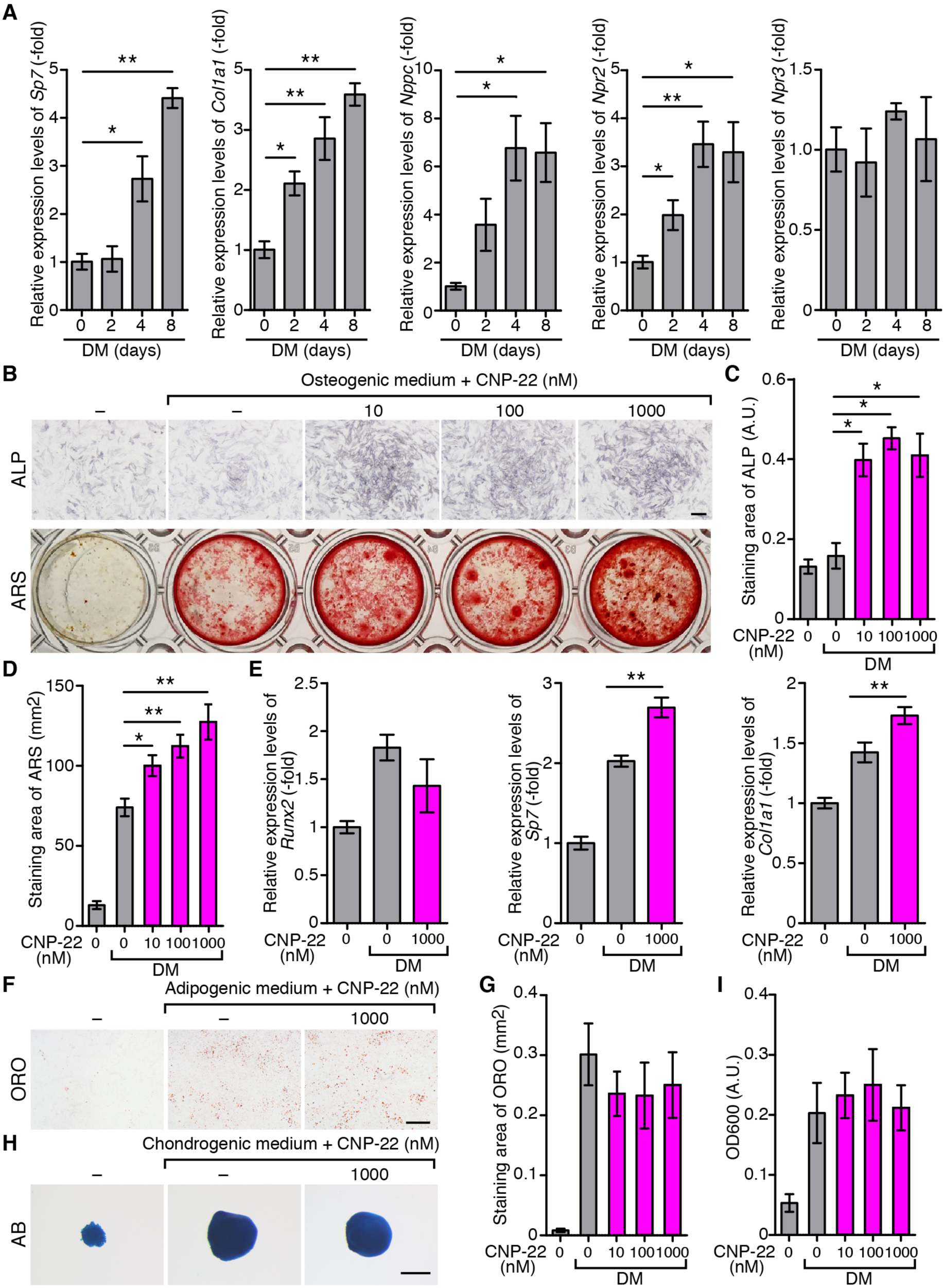
CNP promotes osteogenic differentiation of periosteum-derived cells. (A) Expression analyses of osteogenic marker and CNP-related genes at the days (indicated on x-axis) after the induction with differentiation medium (DM) in periosteum-derived cells (PDCs). Gene expression value of each sample was normalized by each *Gapdh* mRNA value. The relative expression to the value at start point is indicated. (n=3) (B) Representative images of alkaline phosphatase (ALP) activity (upper) and Alizarin red S (ARS) staining (lower) images at 1 day and 8 days of osteogenic differentiation, respectively. (C) Quantified results of ALP-positive areas at 8 days of osteogenic differentiation with or without CNP-22 at the indicated concentration. (n=3) (D) Quantified results of ARS-positive areas with or without indicated concentrations of CNP-22 at 8 days after differentiation. (n=7) (E) qPCR analyses of the expression of osteoblast markers (*Runx2*, *Sp*7, and *Col1a1*) at 4 days of osteogenic differentiation with or without CNP-22 (1000 nM). Data is indicated similarly to (A). (n=4) (F and G) Representative images (F) and quantified results (G) of Oil Red O (ORO) staining images at 8 days of adipogenic differentiation with or without CNP-22. (n=3) (H,I) Representative Alcian Blue (AB) staining (H) of micromass culture and quantified results (I) of AB extracts from samples used in (H) at 14 days of chondrogenic differentiation with or without indicated concentrations of CNP-22. (n=4) Scale bars: B, 0.5 mm; F, 1 mm; H, 50 μm. Data are mean ± s.e.m. ** *P* < 0.01, \**P*<0.05; by One-way ANOVA followed by Turkey-Kramer test.

### OSTN and CNP synergistically augment osteogenic differentiation of PDCs

The previous studies reported that OSTN augments the NP signaling by protecting NPs from NPR3-dependent clearance (Kanai et al., 2017; Kita et al., 2009; Miyazaki et al., 2018; Moffatt et al., 2007; Subbotina et al., 2015). To determine whether OSTN acts as a regulator of the CNP signaling in PDCs, we evaluated the effect of OSTN on CNP-stimulated osteogenic differentiation of PDCs. In the presence of CNP-22 (5 nM), low doses of OSTN (1 and 10 nM) significantly enhanced osteogenic differentiation of PDCs (Fig. 6A upper,B). Furthermore, OSTN significantly promoted osteogenic differentiation without exogenous CNP-22, implying the presence of endogenous CNP-dependent regulation of osteogenesis of PDCs (Fig. 6A lower,B). Moreover, OSTN tended to enhance CNP-induced cGMP production in PDCs (Fig. 6C).

**Fig. 6.**
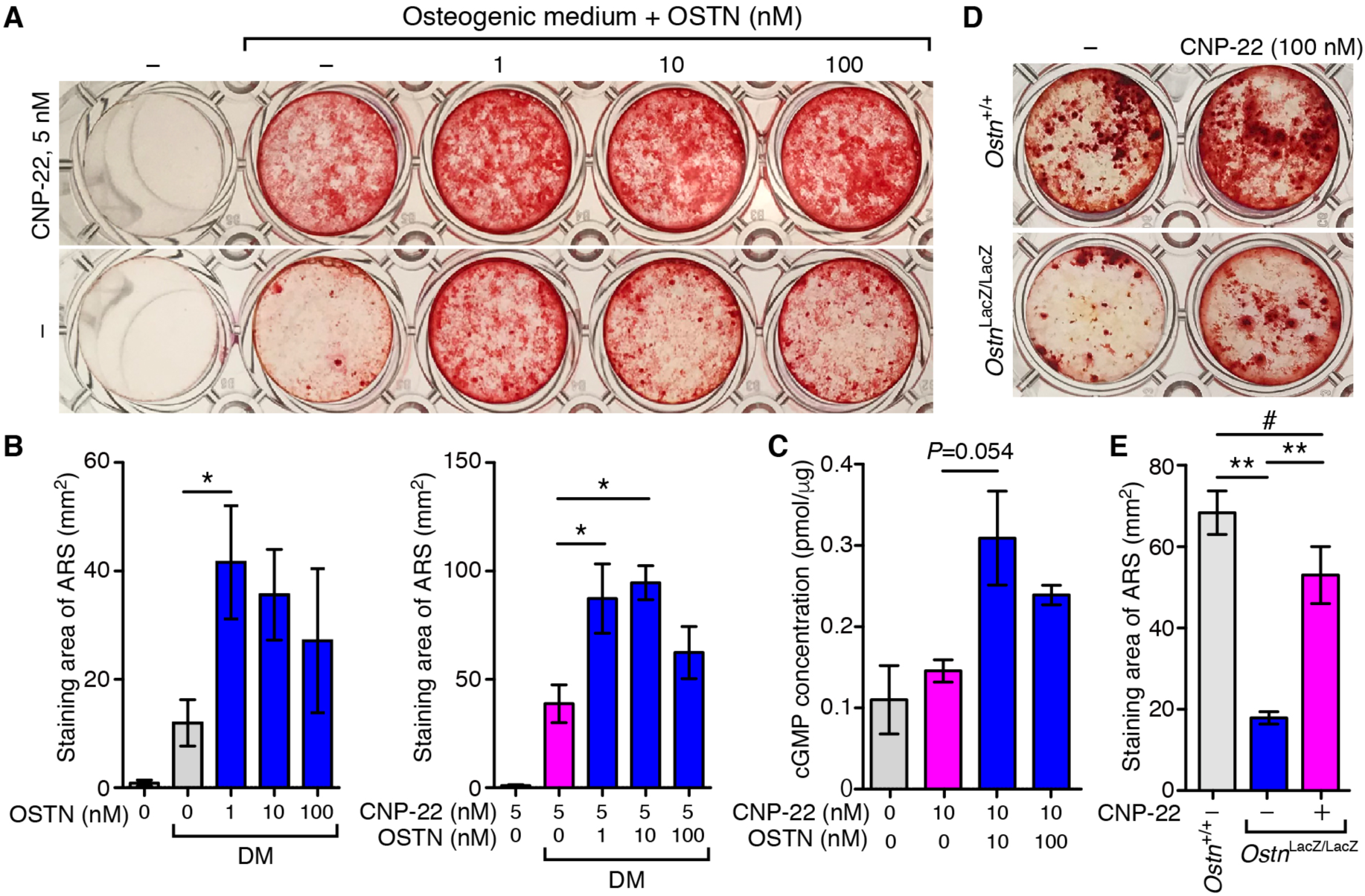
OSTN and CNP synergistically augment osteogenic differentiation of PDCs. (A) Representative ARS images of staining at 5 days of osteogenic differentiation in PDCs with 5 nM (upper) or without (lower) CNP-22 in the combination with the indicated concentrations of OSTN. (B) Quantified result of ARS staining area of images used in (A). (n=4) (C) The effect of indicated concentration of OSTN on cGMP production of PDCs at 15 minutes after the stimulation with low dose of CNP-22 (10 nM). (n=3) (D) Representative ARS staining at 8 days of osteogenic differentiation of PDCs isolated from 8-week-old *Ostn*^LacZ/LacZ^ and *Ostn* ^+/+^ mice. (E) Quantitative analysis of ARS-positive area in the images used in (D). (n=3) Data are mean ± s.e.m. ** *P* < 0.01, * *P*<0.05, # *P*>0.05; by One-way ANOVA followed by Turkey-Kramer test.

To investigate whether CNP-dependent regulation in the periosteum is related to the impaired cortical bone formation of the *Ostn*^LacZ/LacZ^ mice, we isolated PDCs from *Ostn*^LacZ/LacZ^ mice and examined the osteoblast differentiation. The PDCs isolated from *Ostn*^LacZ/LacZ^ mice hardly differentiated into osteoblasts as compared with those from the control mice (Fig. 6D,E). However, this decreased osteogenic differentiation of the *Ostn*^LacZ/LacZ^ PDCs was partially rescued by CNP. These results indicate the reduction of CNP-dependent osteoblast differentiation might account for the reduced cortical bone formation of the *Ostn*^LacZ/LacZ^ mice. Collectively, OSTN promotes osteogenic differentiation of PDCs by augmenting CNP signaling pathway, thereby regulating cortical bone formation.

### OSTN-bound NPR3-expressing cells can differentiate into several lineages

To examine whether OSTN regulates the CNP-GC-B axis via NPR3 in the periosteum, we quantified the expression of *Npr1*, *Npr2*, and *Npr3* using fractionated bone samples: PS, BM, and TB/CB. While the cells obtained from BM highly expressed *Npr1*, those from PS expressed *Npr2* and *Npr3* (Fig. 7A). To characterize NPR3-expressing cells, we isolated cells that bind to OSTN by FACS because OSTN can bind to NPR3 (Fig. 7B). Biotin-tagged OSTN was labeled by Cy3 using streptavidin (SA) and mixed with isolated PDCs. Biotin-tagged OSTN, but not biotin alone, increased Cy3 intensity in about 30% of total PDCs (Fig. 7C). Furthermore, untagged OSTN suppressed the increased number of the cells that bound to OSTN. On the other hand, *Npr3*-deficient PDCs didn’t bind to labeled OSTN. These data suggest that the binding of OSTN to the PDCs depends on NPR3.

**Fig. 7.**
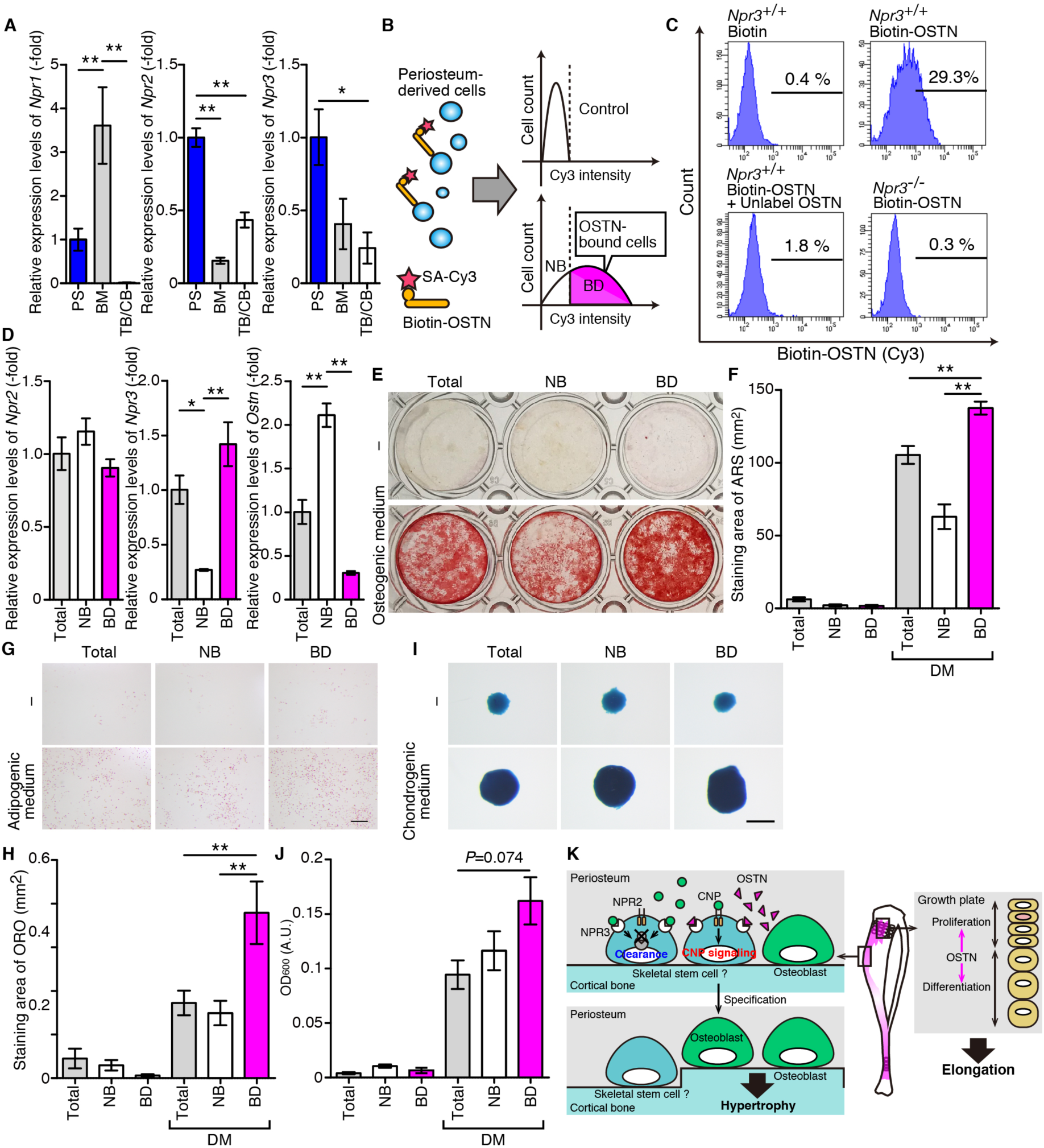
OSTN-bound NPR3-expressing cells can differentiate into several lineages. (A) qPCR analyses of the expression of receptors for natriuretic peptides (*Npr1*, *Npr2*, and *Npr3*) in fractionated bones. Data is indicated similarly to Fig. 1E. (n=4) (B) Schematic representation of strategy for capture the cells targeted by OSTN in the periosteum. Biotin-tagged OSTN was labeled by Cy3 using streptavidin (SA) and mixed with isolated PDCs, followed by analysis using FACS. Cy3 positive and negative cells are collected as OSTN-bound cells (BD) and OSTN non-bound cells (NB), respectively. (C) Representative histograms of FACS analyses under the indicated conditions. x and y axis indicate Cy3 intensity and cell number, respectively. Numbers indicate the percentages of OSTN-binding cells per parent population. (D) qPCR analyses of the expression of *Npr1*, *Npr2*, and *Npr3* in total, NB, and BD fractions of PDCs. Data is indicated similarly to Fig. 1E. (n=3) (E,F) Representative ARS staining images (E) and quantitative analysis (F) of FACS-sorted PDCs cultured in growth medium or differentiation medium (DM) for 8 days. (n=3) (G,H) Representative ORO staining images (G) and quantitative analysis (H) using sorted fraction of PDCs at 8 days after the culture in growth medium or differentiation medium (DM). (n=3) (I,J) Representative AB staining images (I) and quantitative analysis (J) of AB extracts using sorted fraction of PDCs at 14 days after the culture in growth medium or differentiation medium (DM). (n=3) (K) Proposed schematic diagram of OSTN roles in bone development. OSTN promotes endochondral ossification, leading to long-axis growth of bones through possibly enhancing proliferation and maturation of chondrocytes. On the other hand, CNP signaling augmented by OSTN in NPR3-expressing cells leads to specification into osteoblasts, thereby promoting short-axis growth through intramembranous ossification. Data are mean ± s.e.m. ** *P* < 0.01, * *P*<0.05; by One-way ANOVA followed by Turkey-Kramer test.

To further confirm Npr3-dependent binding of OSTN to PDCs, we analyzed gene expression of *Npr3* and *Ostn* in the OSTN-bound or OSTN-unbound PDCs (Fig. 7D). Consistent with the results of FACS analyses, *Npr3* was highly expressed in the OSTN-bound cells. Conversely, *Ostn* was expressed in the OSTN-unbound cells, indicating that OSTN might act in a paracrine manner in the periosteum (Fig. 7D). Because periosteum includes large amount of skeletal stem cells that give rise to osteoblasts, chondrocytes, and adipocytes (van Gastel et al., 2012), we tested whether OSTN-bound cells can differentiate into several lineages. OSTN-bound cells were prone to differentiate into several lineages: osteogenic, adipogenic, and chondrogenic, when they were cultured under appropriate conditions as compared with total and OSTN-unbound cells (Fig. 7E-J). These results suggest that OSTN targets Npr3-expressing cell that might function as skeletal stem cell in the periosteum.

## DISCUSSION

The present study for the first time demonstrates the essential and physiological role for periosteum-derived peptide, OSTN, in the mammalian bone growth. By analyzing OSTN-LacZ reporter gene knock-in mouse, we found *Ostn* expression in the periosteum of a part of long bones as well as flat bones. Furthermore, analyses of OSTN-deficient mice revealed that OSTN is required for both types of long bone growth; endochondral and intramembranous ossification through augmenting CNP signaling.

Periosteum-derived OSTN promotes endochondral ossification involved in long bone growth possibly by promoting the CNP-GC-B signaling pathway. Several lines of studies have suggested that the CNP-GC-B axis plays a pivotal role in endochondral ossification during long bone growth (Chusho et al., 2001; Nakao et al., 2013; Yasoda et al., 2004). We have recently shown that overexpression of OSTN leads to the elongation of bones in both zebrafish and mouse in a CNP-dependent manner (Chiba et al., 2017; Kanai et al., 2017). This study further demonstrates the role of endogenous OSTN in regulation of endochondral ossification during long bone growth. Although our data together with previously reports suggest that OSTN regulates endochondral ossification by potentiating the CNP-GC-B pathway through inhibition of NPR3-mediated degradation of CNP, it remains unclear how OSTN spatially regulates the CNP-GC-B signaling axis. CNP, GC-B, and cGKII have been shown to be expressed in proliferating and prehypertrophic zones of growth plate, while NPR3 is expressed in hypertrophic chondrocytes. However, our analyses on OSTN clearly showed the expression of OSTN in periosteal-osteoblasts, but not in the growth plate. Thus, periosteum-derived OSTN needs to be delivered to the growth plate.

Although there are no penetrating blood vessels into the layers of growth plates, epiphyseal and metaphyseal ends of growth plate are supplied by epiphyseal arteries and metaphyseal arteries, respectively (Ramasamy, 2017). In addition, periosteal blood vessels have been shown to pass across cortical bone and anastomose with the arterioles of the endosteum (Spira and Farin, 1967). Therefore, OSTN released from the periosteal-osteoblasts may be delivered to the growth plate through metaphyseal arteries to spatially regulate the CNP-GC-B signaling. However, further studies are needed to confirm this hypothesis.

OSTN expressed in the periosteum regulates appositional growth as well as endochondral ossification. OSTN-deficient mice exhibited the morphological defects in the cortical bone of tibia. They also showed decreased number of osteoblasts in the periosteum/perichondrium of developing tibiae compared with wild-type mice. It has been shown that bones become wider through the process by which periosteal osteoblasts add mineralized tissues beneath periosteum, i.e. intramembranous ossification (Rauch, 2005). Furthermore, recent study shows that periosteal stem cell-derived osteoblasts promote appositional growth of long bones (Debnath et al., 2018). Therefore, OSTN regulates appositional growth of long bones by inducing osteoblast-dependent intramembranous ossification.

OSTN might facilitate osteogenic differentiation of periosteal skeletal stem cells by enhancing the CNP-GC-B signaling pathway. In this study, we found that osteogenic differentiation of PDCs that contain periosteal skeletal stem cells was synergistically enhanced by OSTN and CNP. More importantly, PDCs derived from OSTN-deficient mice showed less ability to differentiate into osteoblasts than the control. Furthermore, their defects of stem cell differentiation were partially rescued by the stimulation with CNP. Therefore, OSTN and CNP might cooperatively regulate osteogenic differentiation of periosteal stem cells. Indeed, PDCs contained a population of NPR3-expressing cells that have the potential to differentiate into several lineages including adipocytes, chondrocytes, and osteoblasts. Thus, OSTN may promote CNP-mediated osteogenic differentiation of NPR3-expressing multipotent stem cells by preventing the NPR3-mediated clearance of CNP. However, it is important to clarify whether NPR3-expressing cells in the periosteum are identical to periosteal skeletal stem cell involved in intramembranous bone formation.

OSTN may spatiotemporally regulate appositional growth in the periosteum. The expression of CNP and GC-B was abundant in periosteum and increased during osteogenic differentiation of PDCs, suggesting that osteogenic cells became more sensitive to the stimulation of CNP. On the other hand, OSTN was expressed in periosteum-derived osteoblasts and synergistically promoted osteogenic differentiation of PDCs with CNP. Accordingly, OSTN is likely involved in a positive feedback mechanism of osteogenic differentiation in PDCs. The spatiotemporal expression pattern of OSTN demonstrated by OSTN-LacZ knock-in mice suggest the involvement of OSTN in the local regulation of bone growth. In fact, appositional growth is likely to be influenced by age- and location-dependent variables. Radiographic studies using human subjects indicate the rate of appositional growth reaches a peak in adolescence and site-specific within the bone (Baxter-Jones et al., 2011; Gabel et al., 2015). Therefore, OSTN may regulate bone growth as a local cue in the periosteum to attain site-specific appositional growth.

There might be an unexpected function for the CNP-GC-B signaling axis in osteogenic differentiation of periosteal skeletal stem cells required for intramembranous ossification. Previous in vitro studies have shown that CNP stimulates osteogenic differentiation of rat calvaria-derived osteogenic cells and MC3T3-E1 (34-36). In addition, CNP enhances proliferation of human osteoblasts in vitro (Lenz et al., 2010). However, in vivo role of CNP for osteogenic differentiation is more complicated than in vitro. The mice overexpressing CNP under the control of serum amyloid promoter (SAP-CNP Tg mice) exhibit decreased trabecular bone volume and cortical thickness because of increased bone turnover (Kondo et al., 2015). Consistently, bone matrix, serum ALP, and related bone metabolic parameters in urine are increased in *Npr3*^-/-^ mice, suggesting the increased bone metabolism in this mutant (Matsukawa et al., 1999). However, new bone formation and bone strength at the fracture sites are promoted in SAP-CNP Tg mice (Kondo et al., 2015). Furthermore, long bone abnormality (*lbab*/*lbab*), a mutant strain carrying a mutation in the *Nppc* gene, shows decreased bone volume of humerus (Kondo et al., 2012). These studies suggest that CNP signaling might be tightly regulated to maintain proper concentration or might be counter-balanced by CNP-clearing system in the tissue in vivo. Thus, further studies are required to elucidate the precise molecular mechanism underlying CNP signal-mediated bone development, metabolism, and fracture healing.

Our study provides the new insight into the mechanism of osteogenic differentiation from NPR3-positve cells in periosteum. It has been widely accepted that periosteum is a major source of skeletal stem cells that repair bone injury (Neagu et al., 2016). A few studies clearly suggested periosteal skeletal stem cells exhibit distinct properties and extensively respond more favorably to bone injury than bone marrow-derived stem cells (Debnath et al., 2018; Duchamp de Lageneste et al., 2018). Furthermore, the expression of CNP, GC-B, and NPR3 is increased in femurs during the healing process of fractured bone in mice (Kondo et al., 2015). Therefore, CNP-GC-B axis might have promotive effects on fracture healing as well as appositional growth. Thus, increasing the level of OSTN or CNP in blood stream may accelerate the fracture healing process. This will be a subject for future study.

In conclusion, we elucidated here periosteum-derived OSTN promotes bone growth through endochondral ossification and intramembranous ossification during murine skeletal development. Thus, our study not only reveals physiological role of OSTN in skeletal development, but also provide new molecular mechanism of intramembranous ossification for appositional growth in the periosteum.

## MATERIALS AND METHODS

### Antibodies and peptides

Antibodies and peptides used here were purchased as follows: rabbit pAb anti-Osterix (ab22552) from Abcam; Dako EnVision+ System-HRP Labelled Polymer anti-Rabbit (K4002) from Dako. CNP-22 (4229-v) from PEPTIDE INSTITUTE, inc., OSTN and Biotin-tagged OSTN from Toray Research Center, Inc.

### Whole mount LacZ staining and clearing

Detection of LacZ activity was performed using 5-Bromo-4-chloro-3-indolyl β-D-galactopyranoside (X-gal). Animals were anesthetized, perfused with PBS, and fixed with fixative solution containing (0.1 M Na-PO4 buffer [pH7.3], 1 % paraformaldehyde (PFA), 0.05 % glutaraldehyde, 5 mM EGTA, 2 mM MgCl2, and 0.1 % NP-40). The deskinned animals were incubated in staining solution (0.1 M Na-PO4 [pH7.3], 2 mM MgCl2, 0.01 % sodium deoxycholate (DOC), 0.02 % NP-40, 5 mM K-ferricyanide, 5 mM K-ferrocyanide, 20 mM Tris-HCl [pH7.3], and 1 mg/ml X-gal) for overnight at RT. They were washed with PBS and fixed with 4 % PFA/PBS NaN_3_ for 1 hour at 4 °C. Clearing was carried out with CUBIC according to the modified method described previously (Tainaka et al., 2014). Briefly, animals were soaked in 1/2 CUBIC overnight at 37 °C with gentle shaking and transferred to CUBIC for 1 week at 37 °C.

### Micro-CT

Tibiae and femora of mice at 8 weeks were collected, fixed with 4 % PFA/PBS NaN_3_ (Wako) overnight, and stored in 70 % EtOH at 4 °C until analysis. Micro-CT analysis was conducted according to the method described previously (Xu et al., 2016).

### Generation of *Ostn*^LacZ/LacZ^ mouse

A targeting vector that harbors LacZ and neomycin cassette (PG00061_Z_2_D09) in *Ostn* locus was obtained from KOMP. The targeting vector was linearized with NotI and transfected into R1 ES cells by electroporation (Bio-Rad) followed by the culture in the medium containing 250 μg/ml G418 (Wako) for 1 week. G418-resistant clones were selected by genomic PCR and homologous recombination in ES clone was confirmed by genomic PCR or southern blotting. Chimeric mice were backcrossed into C57BL/6J background (F9). To standardize the size of pups, we used the offspring from heterozygous parent and restricted the number of pups to 5 per one mother from P7. We routinely determined the genotypes of offspring by genomic PCR using KOD FX Neo DNA polymerase (TOYOBO) with primers listed in Table S1. Protocols for mice were approved by the Institutional Animal Care and Use Committee of National Cerebral and Cardiovascular Center Research Institute.

### Bone length measurement

Bone lengths of calvaria, femora, tibia, and caudal vertebra were measured by Latheta LCT-200 (Hitachi). The lengths of femora and tibia were calculated by the right and left bone lengths. The lengths of caudal vertebrae (C) were calculated by measuring the total lengths from C1 to C3.

### *In situ* hybridization

Tibiae were immediately fixed with 4 % PFA/PBS NaN_3_ (Wako) for overnight and then soaked into 20 % EDTA (pH7.4) solution to decalcification for 2-3 weeks. Tibiae were infused with 30 % sucrose and followed by freezing in Tissue-Tec OCT compound (Sakura). Ten μm thick sections were cut from blocks and dried at 37 °C for 2 hours. After washing with PBS, slides were treated with 2 μg/ml Proteinase K (QIAGEN) for 5 minutes at room temperature. Subsequent methods were followed as (Watanabe-Takano et al., 2014). The primers for the construction of probes are listed in Table S1. Staining area detected by ALP was measured by ImageJ and calculated the height of hypertrophic chondrocyte layer.

### Tissue preparation and immunohistochemistry

Tibiae were fixed with 4 % paraformaldehyde/PBS NaN_3_ (Wako) overnight. After decalcification with 20 % EDTA solution (pH7.4), tissues were embedded in paraffin wax. For histological examination, tissue sections were cut from paraffin-embedded blocks of tibia in 4 μm thick and stained with hematoxylin and eosin. For histological analysis of cartilage, Alcian Blue staining was carried out according to the method described previously (Kanai et al., 2017). For immunostaining with anti-Osterix antibody, proteolysis-dependent antigen retrieval was needed. The sections were incubated in 1 mg/ml proteinase K in PBS at room temperature for 5 minutes. The sections were permeabilized with 0.3 % TritonX-100 in PBS NaN_3_. After blocking with blocking buffer (5 % normal goat serum in 1*×* PBS NaN_3_), sections were immunostained with anti-Osterix antibody diluted (1/600) in the buffer containing 3% BSA/ PBS NaN_3_ overnight at 4 °C. Detection was carried out with HRP-conjugated secondary antibody followed by DAB staining (Vector Labs #SK-4001). The sections immunostained with anti-Osterix antibody were quantitatively analyzed by ImageJ. The region of 655 μm *×* 300 μm was extracted and converted image to 8-bit grayscale.

### Preparation of periosteum-derived cells

Periosteum-derived cells were collected according to the modified protocol described previously (van Gastel et al., 2012). Briefly, tibiae were dissected and immediately soaked into PBS. To eliminate the contamination from perichondrium, both ends of tibiae were coated with 4 % low-melting agarose. Tibiae were incubated with isolating medium consist of *α*MEM (GIBCO), 3 mg/ml, collagenase type II (Worthington #LS004176), Dispase (GIBCO #17105-041), GlutaMAX^TM^ (GIBCO), and antibiotics (FUJIFILM) for 10 minutes at 37 °C. The samples were vortex for 10 seconds and this first supernatant was discarded. Again, the samples were incubated with new isolating buffer for 60 minutes at 37 °C. The reaction was stopped with *α*MEM containing 10 % FBS, antibiotics, GlutaMAX^TM^ and filtered with 70 μm cell strainer (FALCON). The cells were collected by centrifuged, suspended with growth medium, and incubated at 37 °C in 5% CO_2_ for 4 days. After the culture for 4 days, the cells were passaged and grown for 3 days. To induce osteogenic and adipogenic differentiation, cells were plated in 24 well at 2.5 × 10^4^ cells per well and grown in normal medium for 48 hours. Osteogenic medium contains 10 mM *β*-glycerophosphate, 50 μg/ml ascorbic acid, and adipogenic medium is supplemented with 500 μM isobutylmethylxanthine (IBMX), 2.5 μM dexamethasone (Dex), and 10 μg/ml insulin. For chondrogenic differentiation, 2.5 × 10^5^ cells were plated in a 96-well U-bottom suspension culture plate. After the spontaneous formation of spheroid for 48 hours, medium was exchanged with differentiation medium (PromoCell) with or without CNP-22. Differentiation medium was changed every other day for 14 days.

### Sorting of LacZ-positive cells

Sorting of the cells expressing LacZ was conducted using FluoReporter^TM^ lacZ Flow Cytometry Kit (Invitrogen^TM^). Periosteum-derived cells were isolated according to the above-mentioned method and reacted with fluorescein di-V-galactoside (FDG). The cells showing high FITC intensity was sorted by FACS Aria III (BD). Sorted cells were centrifuged and lysed with TRIzol RNA Isolation Reagents (Thermo Fisher Scientific).

### Estimation of Osterix (+) cells by FACS

Periosteum-derived cells were prepared from mice at the age of P11 according to the above-mentioned method. To eliminate red blood cells, cells were treated with hemolytic reagent and then fixed with 4 % paraformaldehyde/PBS NaN_3_ for overnight at 4 °C. Cells were permeabilized with 0.1 % tritonX-100//PBS NaN_3_ for 5 minutes and incubated for 30 minutes on ice in 1 % BSA/ PBS NaN_3_. Cells were stained with anti-Osterix antibody (1/1200) for 30 minutes on ice and then detected by A488-conjugated secondary antibody.

### Determination of lineage-specific cell differentiation

Cell differentiation was determined by the method described previously (Watanabe-Takano et al., 2010). Mineralization was detected by Alizarin Red S staining. Briefly, cells were fixed with 4 % PFA/PBS NaN_3_ and washed with DDW for several times. Then, the staining solution was added into dishes, and the cells were incubated for 5 minutes at RT. Samples were washed three times with DDW and dried. Adipocytes were assessed by Oil Red O staining. Formalin-fixed cells were washed with DDW and incubated in 0.3 % Oil Red O/60 % isopropanol for 30 minutes at RT. Stained samples were washed with DDW and observed by stereomicroscopy (OLYMPUS). To quantify the differentiation states of samples, stained area of images was measured by ImageJ. For the detection of chondrogenic differentiation, cells were fixed with fixative solution containing 30 % ethanol, 0.4 % PFA, and 4 % acetic acid for 15 minutes at RT. Then, the cells were stained with Alcian Blue solution for overnight at 37 °C and extensively washed with DDW. For quantification, Alcian Blue was extracted by 6 M guanidine hydrochloride solution and measured by optical density at 600 nm (O.D. 600) using spectrophotometer (Bio-Rad).

### OSTN binding assay

Biotin-tagged OSTN was labeled by Cy3-streptavidin (Sigma). Equal volume of 25 μM biotin-tagged OSTN and 0.1 mg/ml Cy3-streptavidin were incubated with on ice for 15 minutes at 4 °C. Cell suspension was incubated with labeled OSTN for 30 minutes at 37 °C. To detect specific binding between OSTN and bound cells, an excess of biotin free OSTN was added before the addition of labeled OSTN. To eliminate the nonspecific binding, the cells were extensively washed with PBS and suspended with 5 % FBS/PBS. Analysis was conducted by BD FACSAria^TM^ III (BD Bioscience).

### RNA preparation and qPCR

Bones were homogenized by physcotron (nition) and isolated total RNA using TRIzol RNA Isolation Reagents (Thermo Fisher Scientific). Purification was carried out RNeasy Plus Kit (QIAGEN) and reverse transcription of mRNA was performed using the SuperScript^TM^ III

Reverse Transcriptase (Thermo Fisher Scientific) according to the company’s protocol. Quantitative PCR was performed on Mastercycler realplex^4^ (Eppendorf) using KOD SYBR® qPCR Mix (TOYOBO) and specific primers. Primers used for real-time PCR were listed in Table S1.

### Morphometric analysis

For morphometric analysis of bones, 10 μl/g calcein solution that contains 2.5 mg/ml calcein, 0.15 M NaCl, 2 % NaHCO3 was administered intraperitoneally at 2 and 5 days before sacrifice. Tibiae were isolated, immediately soaked in 4 % PFA/PBS NaN_3_ overnight and stored in 70 % EtOH at 4 °C until analysis. Morphometric parameters of the osteoblasts were calculated (Niigata Bone Science Institute).

### Analysis of proliferating chondrocytes

For labeling of proliferating chondrocytes, 10 mM EdU diluted with 10 % DMSO/PBS was injected intraperitoneally (10 μl/g). The mice injected with EdU were incubated for 4 hours and dissected. They were immediately fixed with 4 % PFA/PBS NaN_3_, transferred into 20 % EDTA (pH7.4) about 3 weeks, and embedded in paraffin. EdU labeled cells were visualized with Click-iT® EdU Alexa Fluor® 488 Imaging Kit according to manufacturer’s protocol.

### Measurement of cGMP activity

Confluent PDCs in 12 well plates were incubated with serum free *α*MEM containing 250 μM IBMX for 10 minutes before the stimulation. Stimulation with CNP-22 and OSTN was performed for 15 minutes at 37 °C. cGMP concentration was determined using Cyclic GMP ELISA Kit (Cayman chemical) according the manufacture’s protocol.

### Statistics

All results represent mean ± SEM. Statistical analysis was performed by either 2-tailed Student’s *t* test or 1-way ANOVA followed by Tukey-Kramer test. Statistical significance was defined as *P* <0.05.

## Author contributions

H.W.-T. and N.M. designed the experiments and wrote the paper. H.W.-T. performed the almost analysis. A.C. and H.O. helped with the FACS and μCT analysis, respectively. Other coauthors contributed to the discussion and reviewed the manuscript.

## Acknowledgments

We appreciate Dr. Nobuyuki Takahashi for providing Npr3 KO mice. This work was supported by Grants-in-Aid for Young Scientists (A) (15H05646 to H.W.-T.), for Scientific Research on Innovative Areas (18H04994 to H.W.-T.), and for Scientific Research (C) (18K09050 to H.W.-T.) from Japan Society for the Promotion of Science (JSPS), AMED-CREST (Japan Agency for Medical Research and Development, Core Research for Evolutional Science and Technology) (JP17gm0610010 to N.M.); and grants from the Uehara Memorial Foundation (to H.W.-T), the Mochida Memorial Foundation (to H.W.-T), Meiji Yasuda Life Foundation of and health and welfare (to H.W.-T), and the Okinaka Memorial Institute for Medical Research (to H.W.-T).

---

**Fig. S1.**
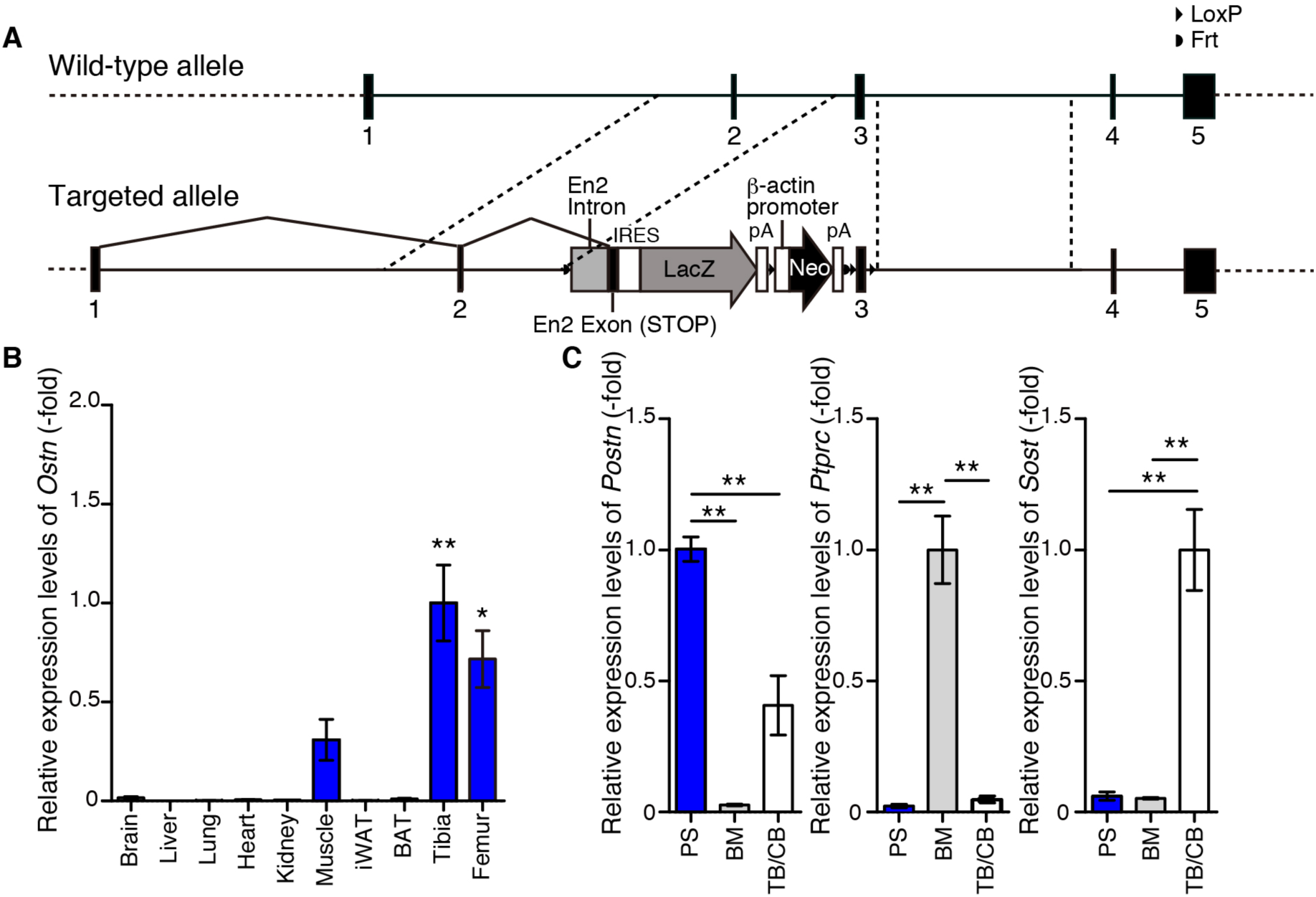
OSTN is expressed in periosteal osteoblasts during bone development. (A) Schematic representation of the targeting method using a vector obtained from KOMP. The targeted allele carries a LacZ reporter gene driven by *Ostn* promoter followed by a Neomycin selection marker induced by *β*-actin promoter. (B) *Ostn* mRNA expression levels in various tissues of ICR male mice at the age of 8 weeks. Gene expression value of each sample is normalized by each 18S rRNA value. The values represent fold change in expression levels relative to that of tibia. (n=4) (C) Validation of fractionated bone samples by analyzing of the expression of specific genes (*Postn*, *Ptprc*, and *Sost)* for each fraction. (n=4) Data are indicated similarly to Fig. 1E. Data are mean ± s.e.m. ** *P* < 0.01, * *P*<0.05; by One-way ANOVA followed by Turkey-Kramer test.

**Fig. S2.**
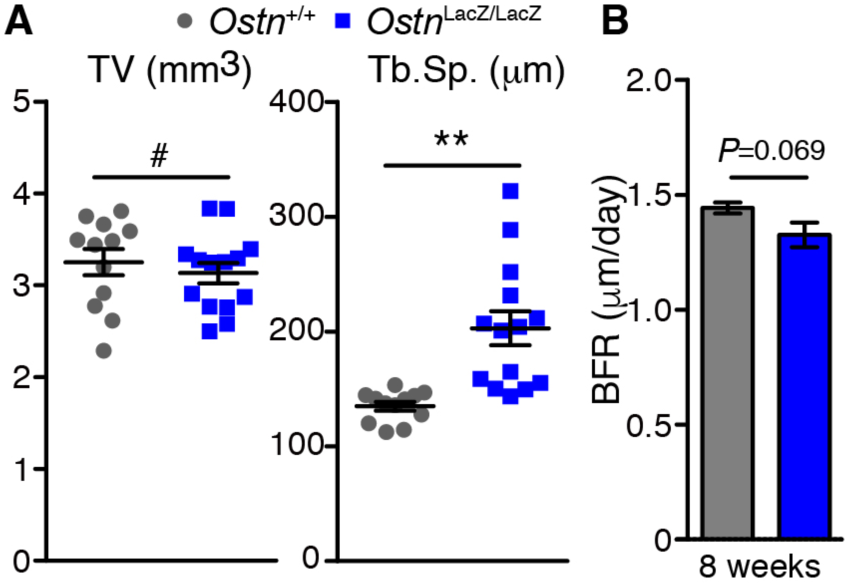
Impairment of growth plate growth and trabecular bone in OSTN-deficient mice. (A) Measurement of trabecular bone parameters: total volume (TV), trabecular space (Tb.Sp.) (n=12-14) (B) Bone forming rate (BFR) assessed by bone morphometric analysis of male 8-week-old *Ostn*^LacZ/LacZ^ and *Ostn*^+/+^ are shown. (n=4), ** *P* < 0.01, #*P* >0.05 by Student’s t test.

**Table S1.**
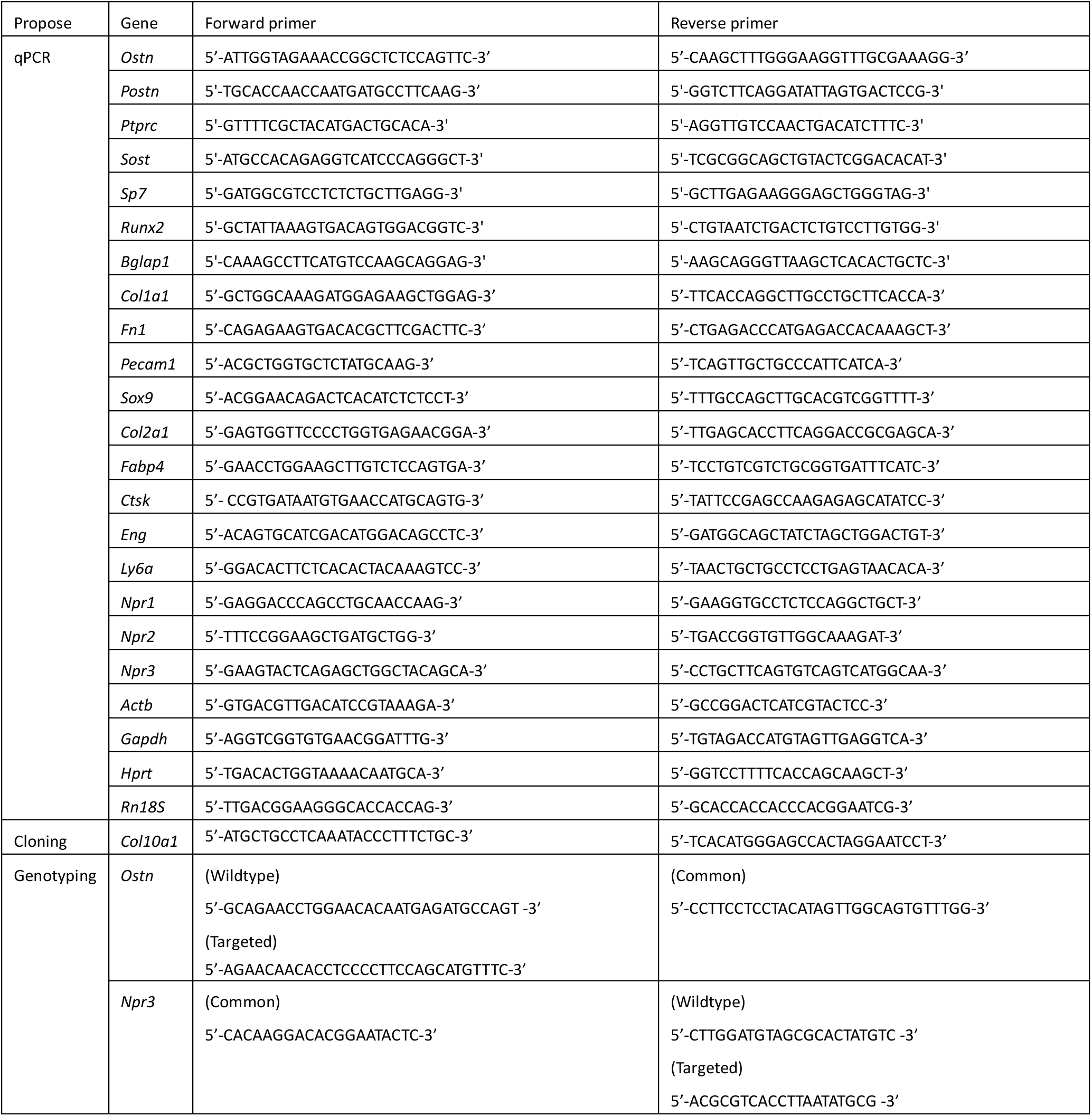
List of primers.

